## Supplemental figures for "β-actin dependent chromatin remodeling mediates compartment level changes in 3D genome architecture"

### SUPPLEMENTARY FIGURES

**Supplementary Figure S1:** ATAC and ChIP signal plotted in 10kb region surrounding center of enhancers and TSSs showing more than two-fold change in ATAC-signal FDR<0.05

**Supplementary Figure S2:** Top 20 GO-Terms associated with A) Genes downregulated by two-fold or more in KO cells with FDR<0.05 B) Genes upregulated by two-fold or more in KO cells with FDR<0.05 C) Genes overlapping EZH2 peaks gaining two-fold or more ChIP-Seq reads in KO cells D) Androgen-Induced genes reported to be upregulated by EZH2 (Zhao et al. Genome Res. 2012 Feb;22(2):322-31) E) TSSs linked to enhancers showing more than two-fold increase in ATAC-signal in KO cells FDR<0.05 F) Promoters showing more than two-fold decrease in ATAC-signal in KO cells FDR<0.05 G) Promoters showing more than two-fold decrease in ATAC-signal in KO cells FDR<0.05 H) TSSs linked to enhancers showing more than two-fold decrease in ATAC-signal FDR<0.05

**Supplementary Figure S3:** Density plots showing average signal intensities (top) and heatmaps displaying scaled read densities (bottom) for ATAC (blue), BRG1 (red), EZH2 (blue), H3K9me3 (green) and H3K27me3 (yellow) in regions ± 5 kb of 1000 random ATAC-Seq peaks. Plots are sorted by WT ATAC signal Scale bar shows normalized RPKM

**Supplementary Figure S4:** A) Normalized counts per million reads for EZH2, BRG1, and REST for wild-type, β-actin knockout and β-actin heterozygous MEFs B) Genome-wide pairwise spearman correlation heatmap of all epigenetic marks in KO (top left triangle) and WT (bottom right triangle) cells

**Supplementary Figure S5:** Pairwise spearman correlation heatmap based on aligned and filtered HiC reads (processed with HiCUP) showing correlation between replicates

**Supplementary Figure S6:** A) ATAC-tracks showing normalized signal (RPKM) at the promoters of select A to B and B to A switching genes B) Top 20 GO-Terms associated with protein-coding genes switching from to A to B (left) or B to A (right) compartments

**Supplementary Figure S7:** A) Average insulation scores of all TAD boundaries B) Average TAD size for all samples. Significance calculated based on Kruskal-Wallis test with Dunn's multiple comparison C) Average signal in RPKM in 200kb region surrounding center of TAD boundaries for i) BRG1 ii)EZH2 iii)H3K9Me3 iv) H3K27Me3 v)ATAC D) ATAC signal in RPKM in 10kb region surrounding ENCODE CTCF Peaks

**Supplementary Figure S8:** A) Average GC (left) and CpG island (right) percentage for 500kb bins belonging to different compartments. Significance calculated using Kruskal Wallis test with Dunns multiple comparisons B) Average GC (left) and CpG island (right) percentage in EZH2 peaks upregulated in KO cells vs all other EZH2 peaks. Significane calculated using two tailed Mann Whitney test C) Average normalized counts (RPKM) per 500kb bin in each compartment type for i) ATAC, ii) H3K9Me3 iii) BRG1 in WT cells

**Supplementary Figure S9:** Scatterplots showing relationship between differentt epigenetic marks and compartment switching using a bin size of A) 50kb and B) 250kb. Scatterplots show KO over WT Log2FC in normalized counts for each genomic bin on the y-axis and difference between KO and WT PC-1 value for each genomic bin on the x-axis. R-Squared and line of best fit for switching bins only (A to B and B to A) shown in black, R-Squared and line of best fit for stable bins only (A and B) shown in grey.

**Supplementary Figure S10:** Pairwise spearman correlation heatmap based on aligned and filtered reads showing correlation between replicates for A) EZH2 ChIP-Seq B) ATAC-Seq

### Figure S1

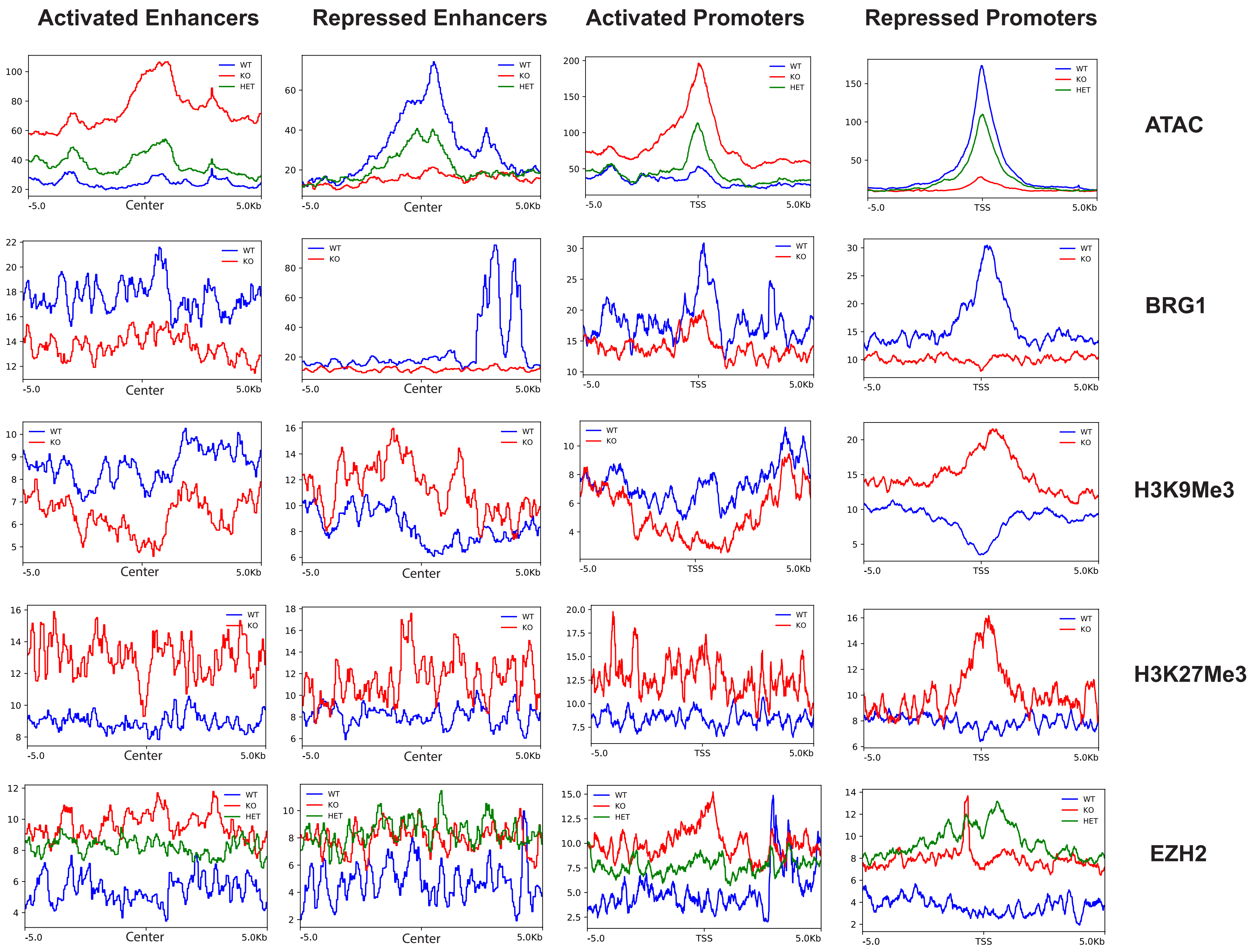

Activated Enhancers
Repressed Enhancers
Activated Promoters
Repressed Promoters
ATAC
BRG1
H3K9Me3
H3K27Me3
EZH2
WT
KO
HET
-5.0
Center
TSS
5.0Kb

### Figure S2

#### GO-TERM ANALYSIS

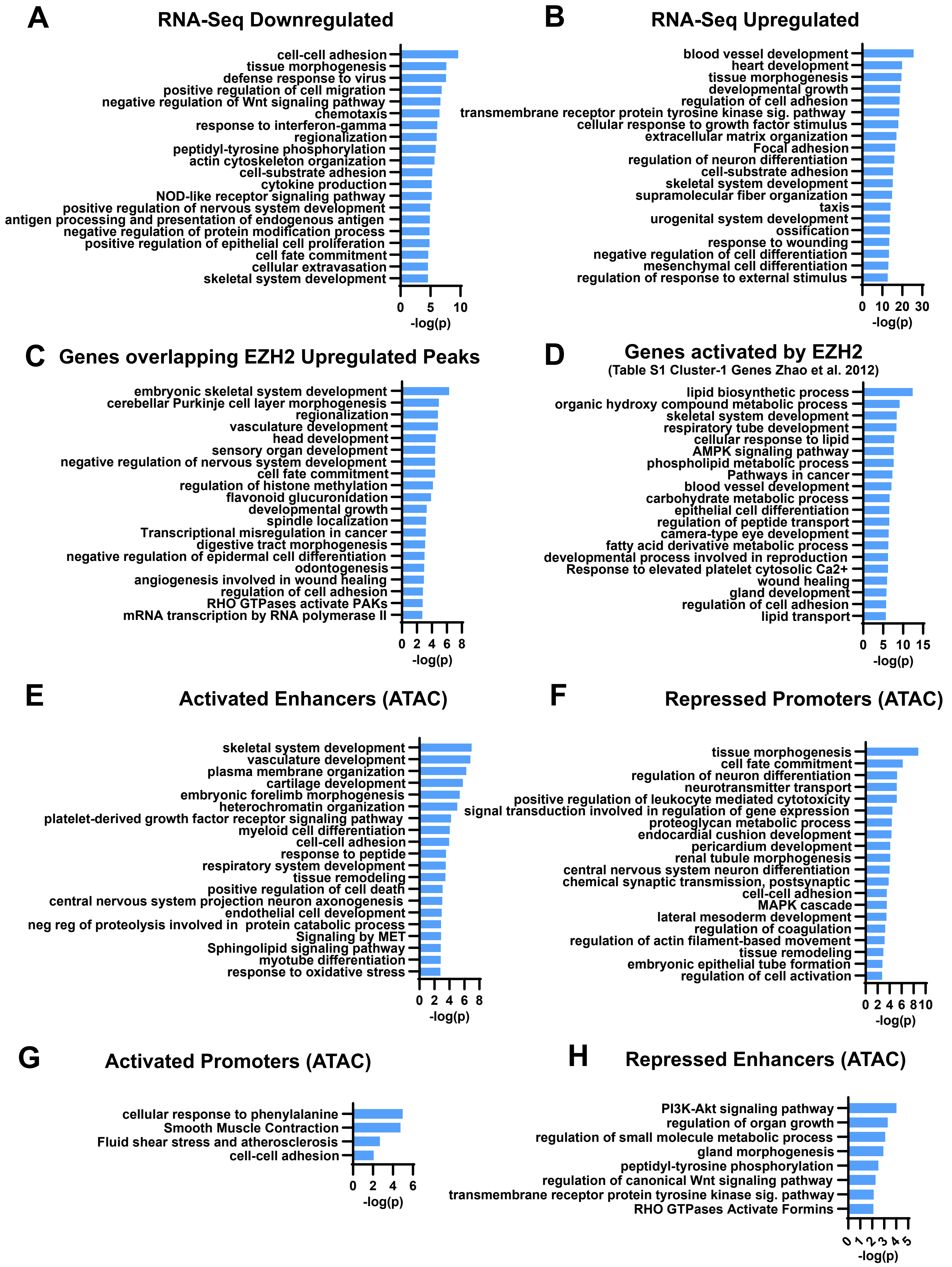

### Figure S3

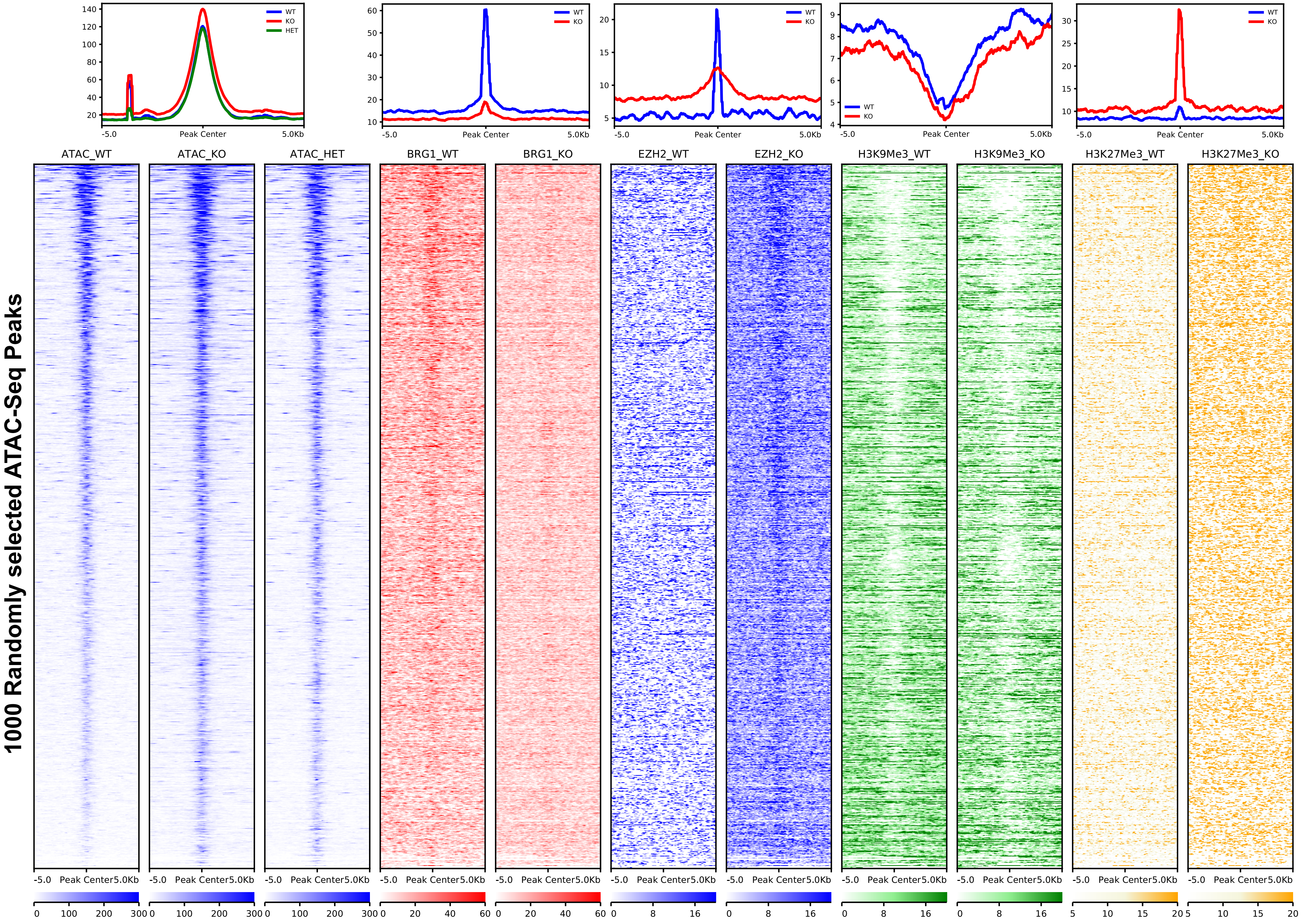
WT
KO
HET
Peak Center
-5.0
5.0Kb
1000 Randomly selected ATAC-Seq Peaks
ATAC_WT
ATAC_KO
ATAC_HET
BRG1_WT
BRG1_KO
EZH2_WT
EZH2_KO
H3K9Me3_WT
H3K9Me3_KO
H3K27Me3_WT
H3K27Me3_KO

### Figure S4

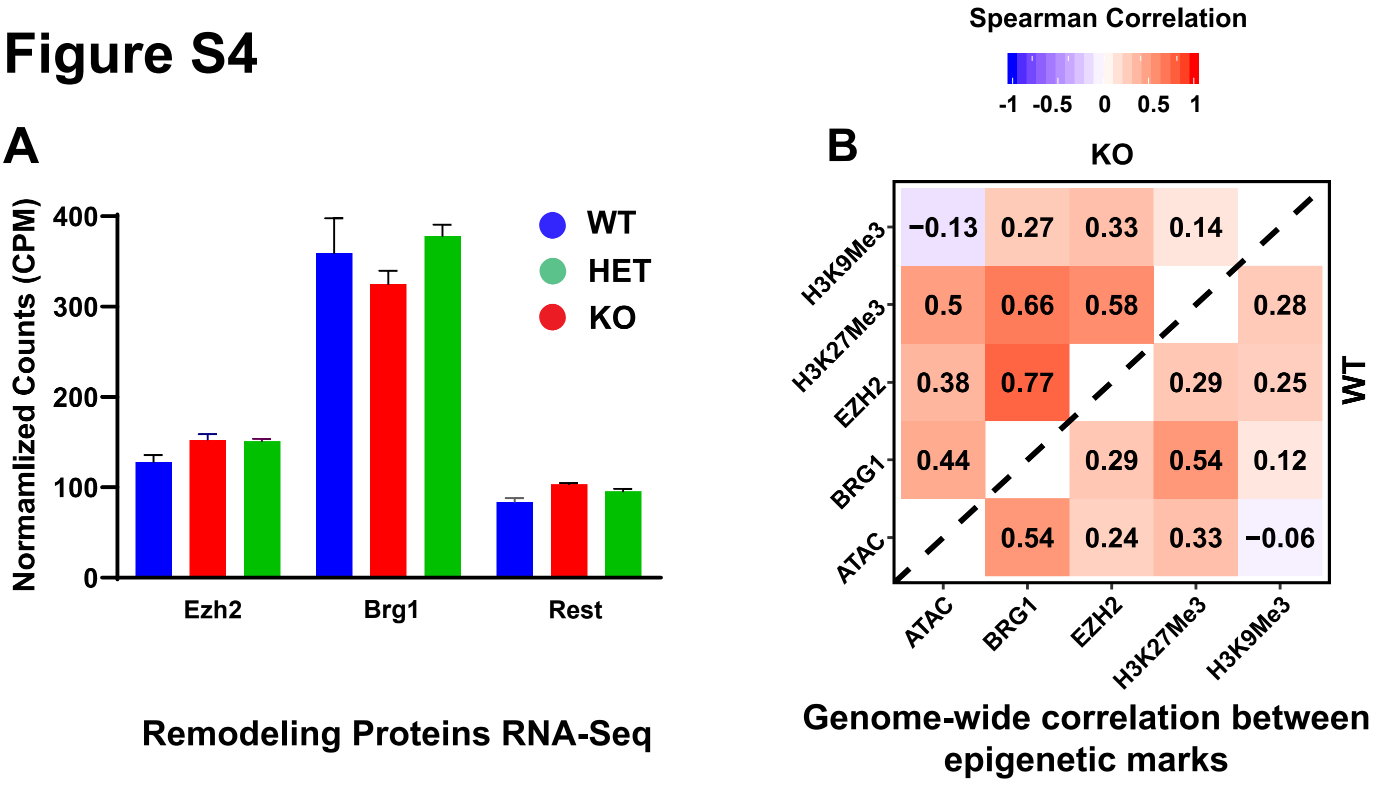


### Figure S5

HiC-Seq Spearman correlation between replicates based on bam alignments

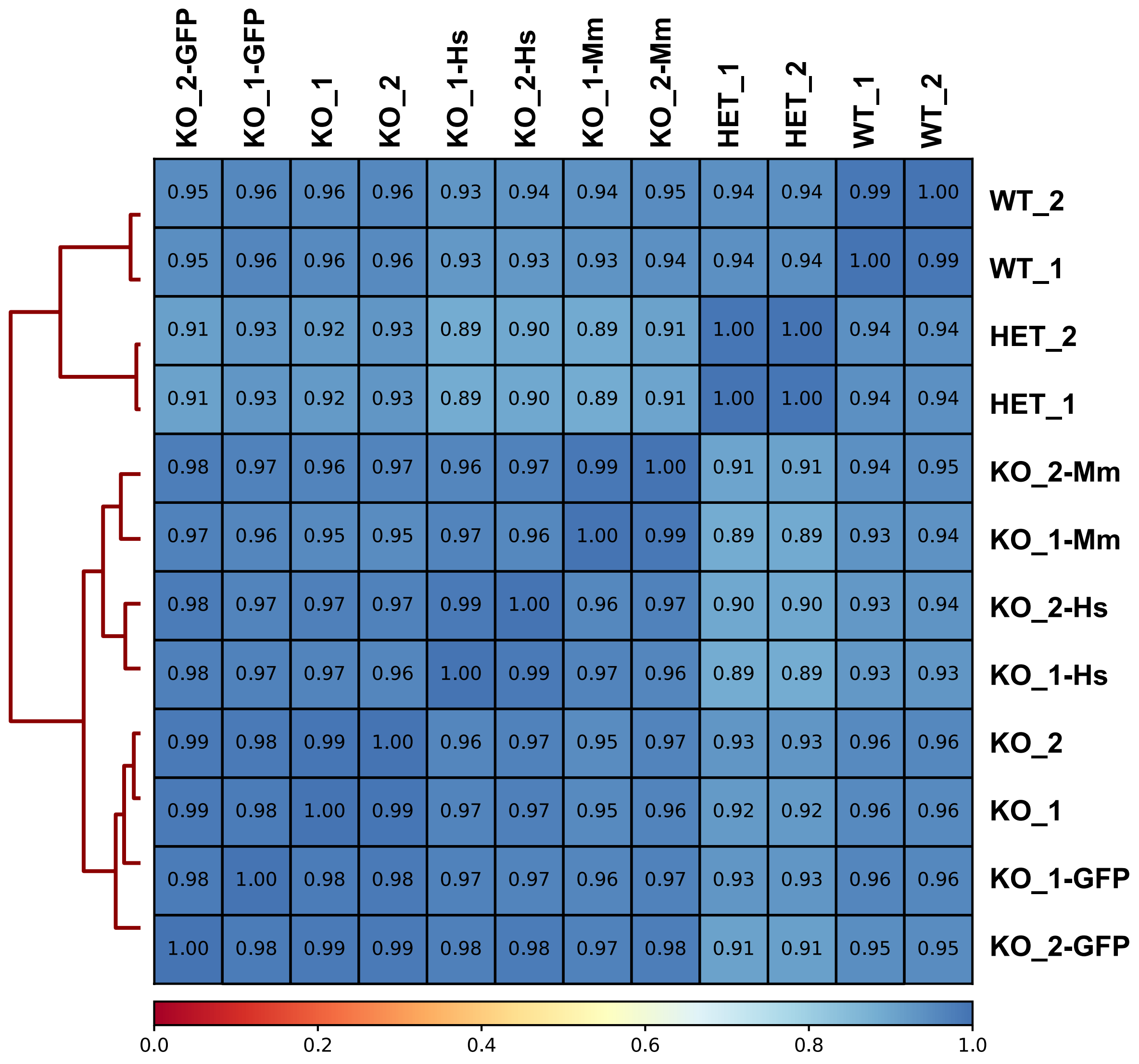

### Figure S6

## A

##### ATAC tracks of selected gene promoters within switching compartments

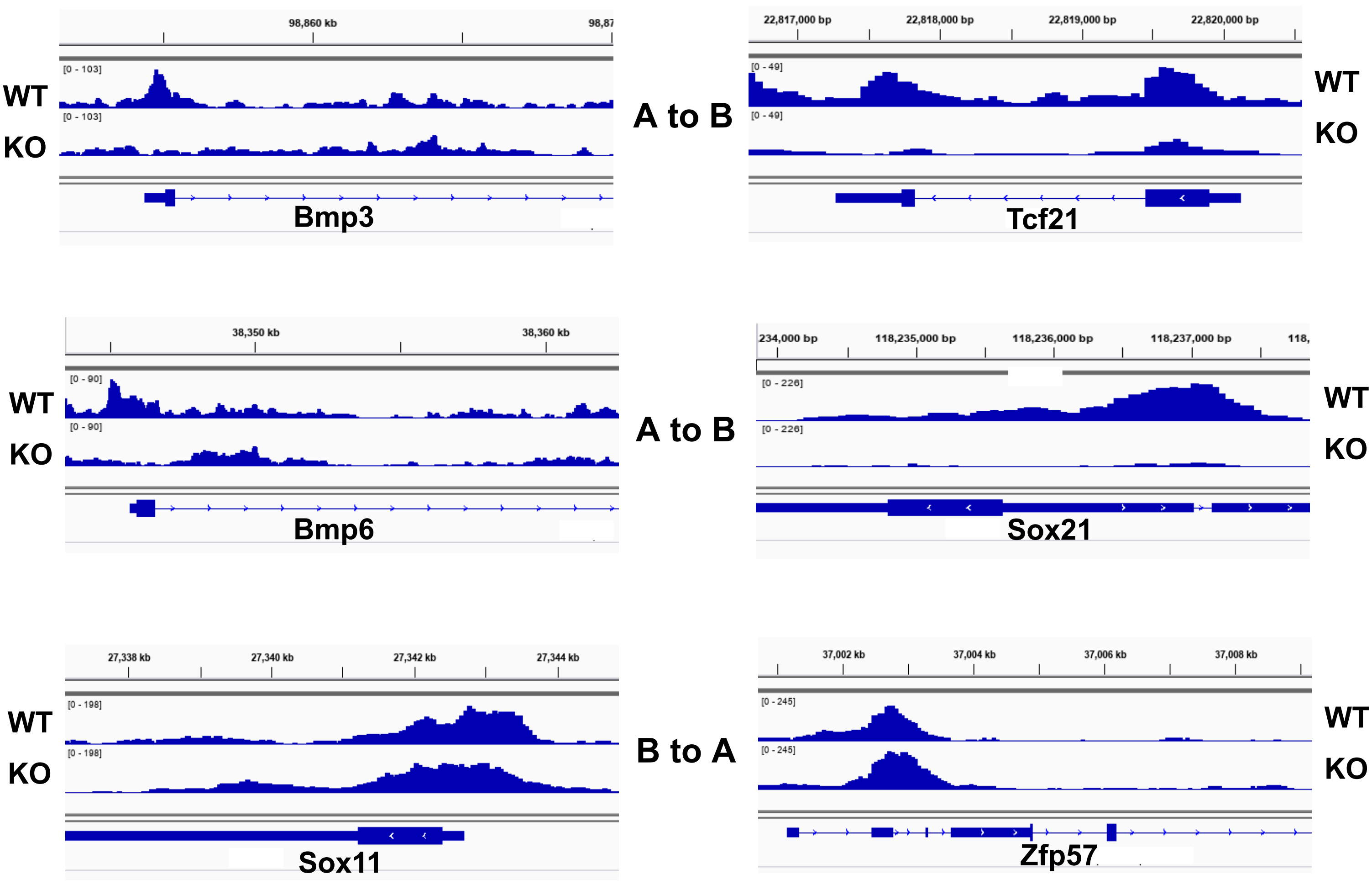


##### GO-Term Analysis

## B

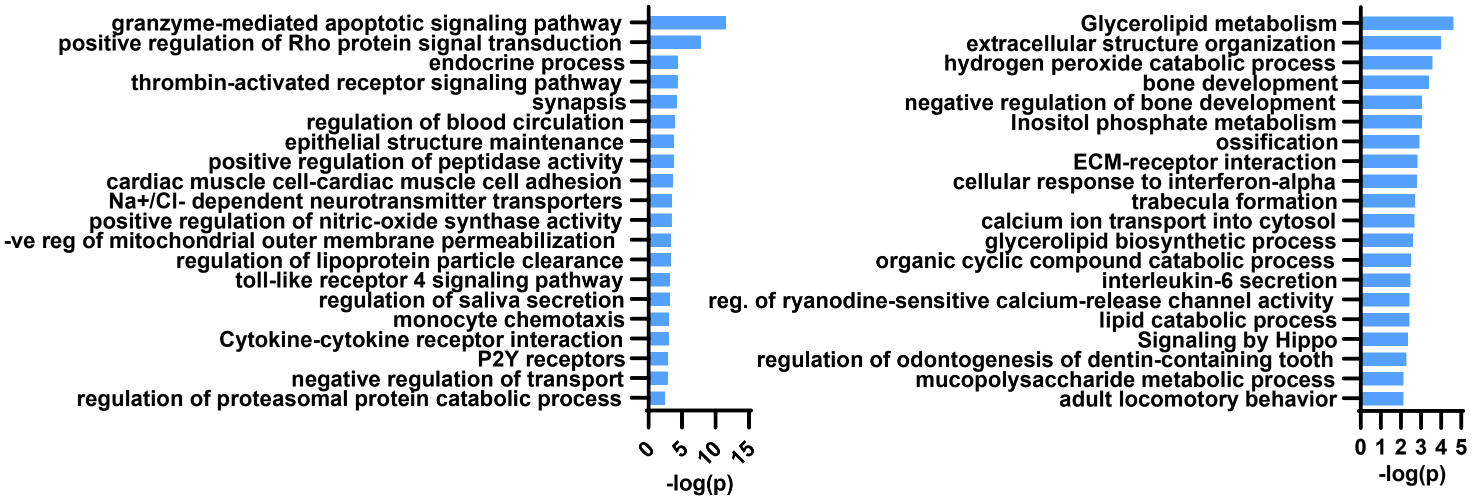

### Figure S7

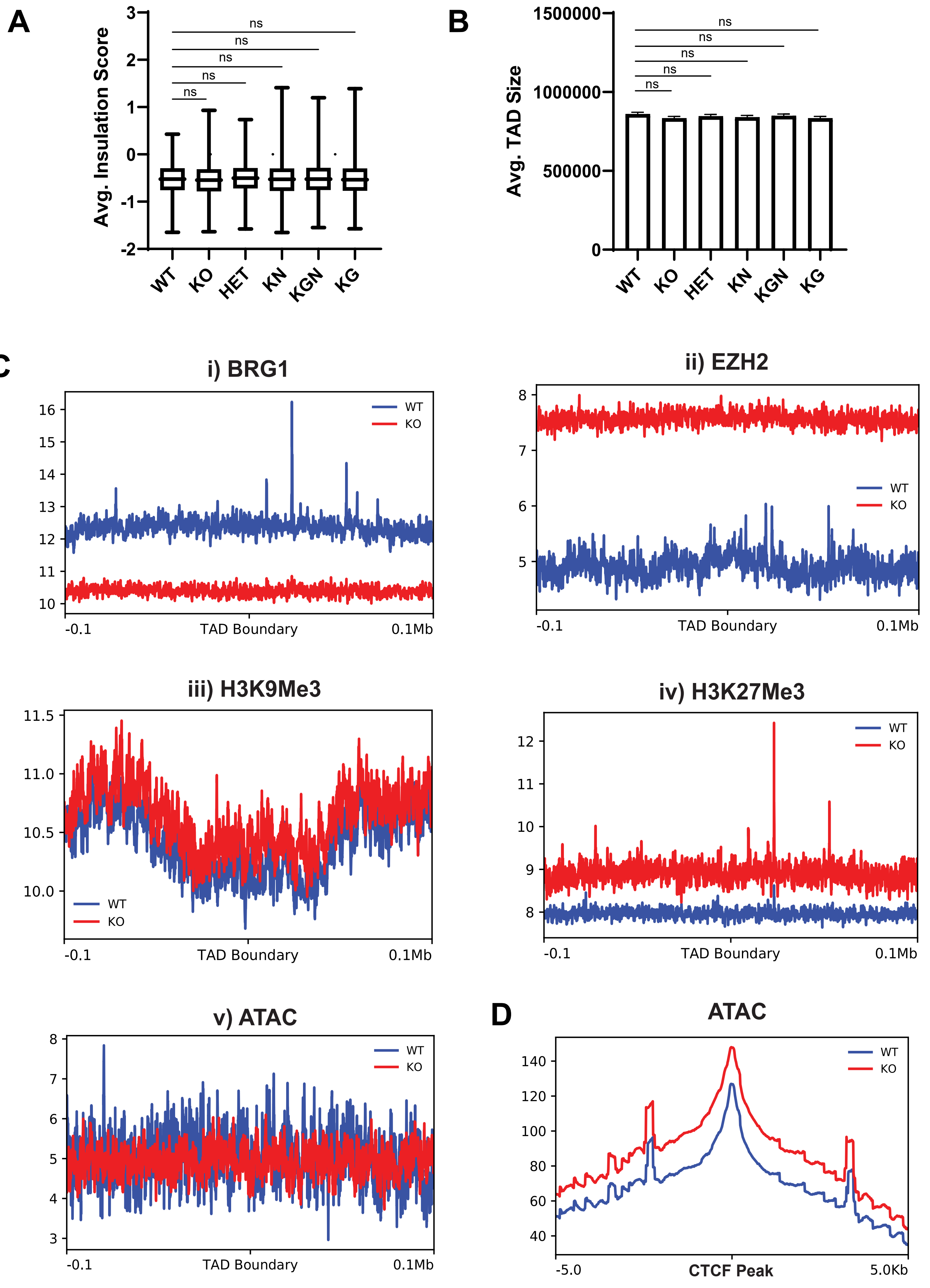
A
Avg. Insulation Score
ns
ns
ns
ns
ns
WT
KO
HET
KN
KGN
KG
B
Avg. TAD Size
1500000
1000000
500000
0
C
i) BRG1
ii) EZH2
iii) H3K9Me3
iv) H3K27Me3
v) ATAC
WT
KO
-0.1
TAD Boundary
0.1Mb
D
ATAC
-5.0
CTCF Peak
5.0Kb

### Figure S8

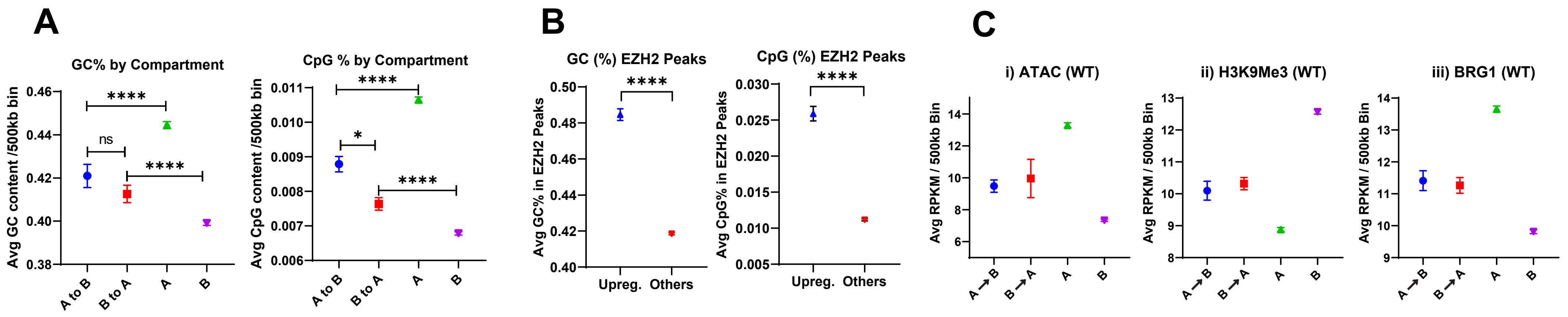


### Figure S9

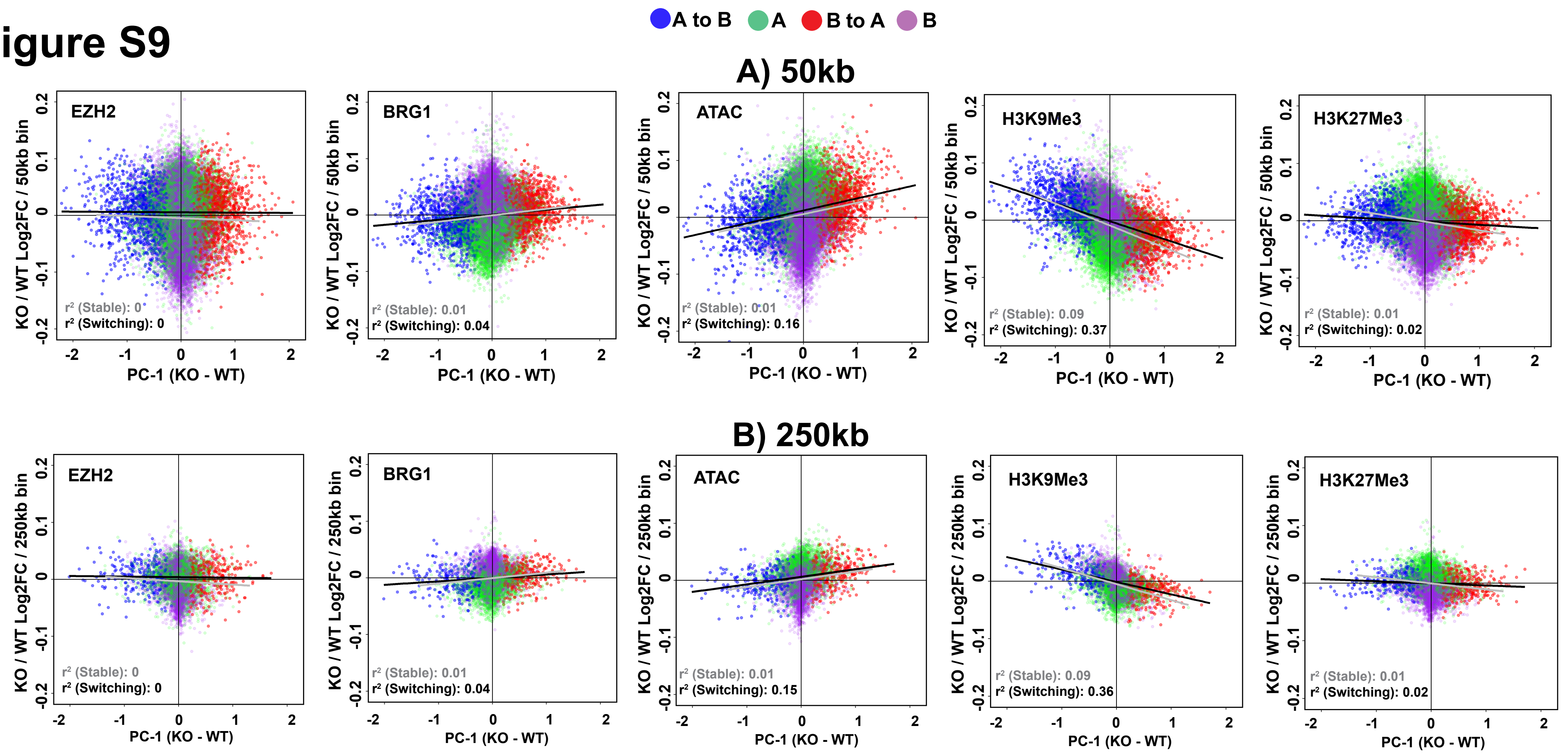

### Figure S10

**EZH2 Chip-Seq Spearman correlation between replicates**

**A**

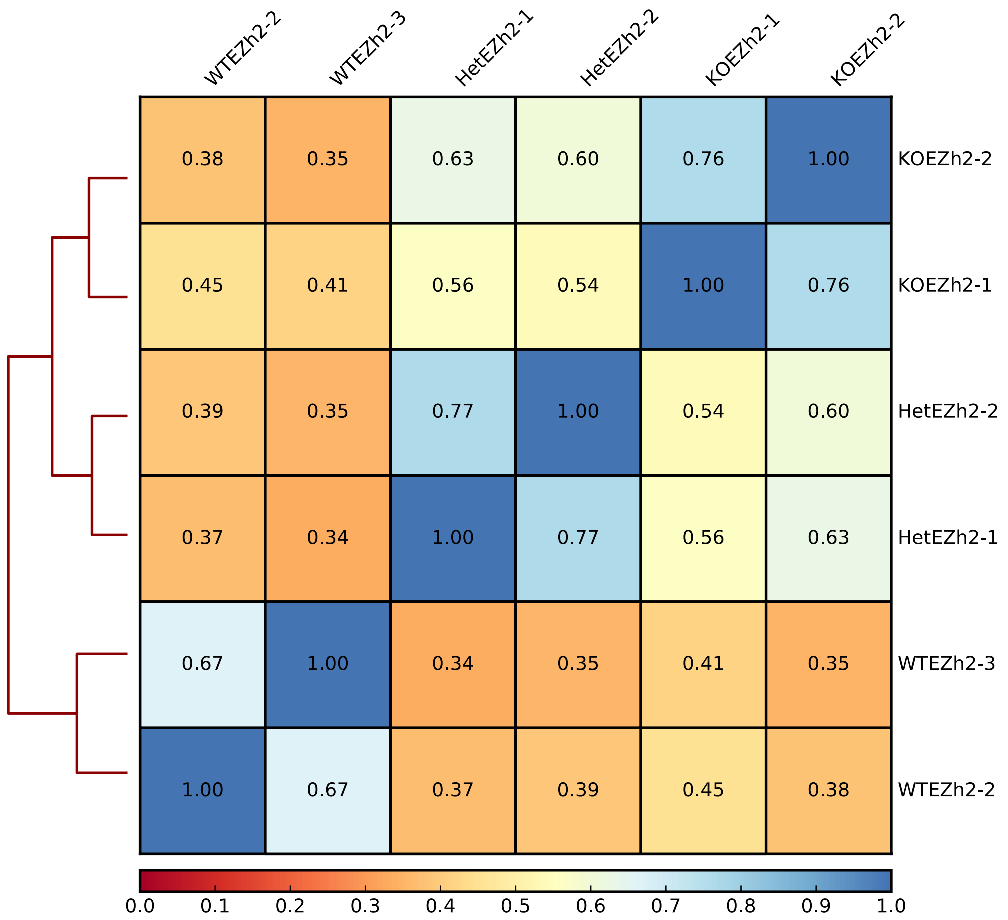


**ATAC-Seq Spearman correlation between replicates**

**B**

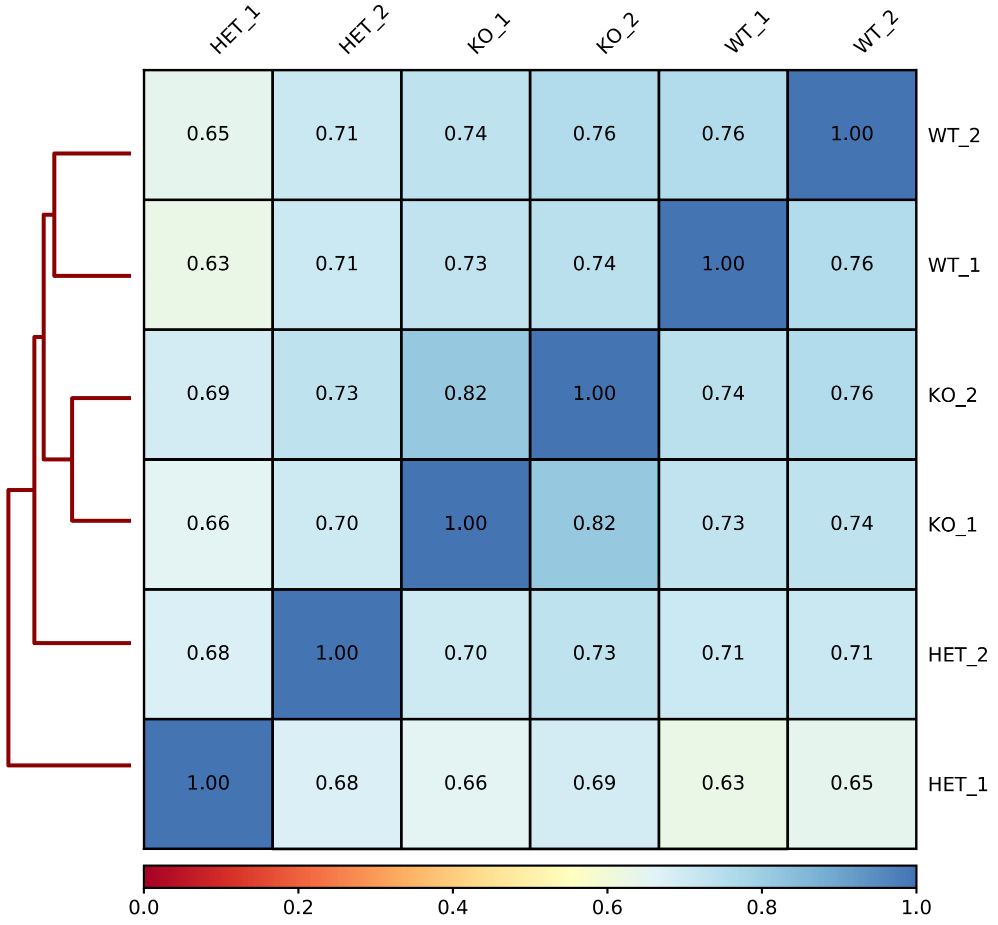
